# Evaluation of metagenomic, 16S rRNA gene and ultra-plexed PCR-based sequencing approaches for profiling antimicrobial resistance gene and bacterial taxonomic composition of polymicrobial samples

**DOI:** 10.1101/2022.05.12.491637

**Authors:** KK Chau, W Matlock, B Constantinides, S Lipworth, L Newbold, H Tipper, T Goodall, H Brett, J Hughes, DW Crook, DW Eyre, DS Read, AS Walker, N Stoesser

**Author notes:** Corresponding author: Kevin K. Chau, Nuffield Department of Medicine, University of Oxford, John Radcliffe Hospital, Headley Way, Headington, OX3 9DU. Nuffield Department of Medicine, University of Oxford, John Radcliffe Hospital, Headley Way, Headington, OX3 9DU.

## Abstract

**Background:** Shotgun metagenomic sequencing is increasingly popular in taxonomic and resistome-profiling of polymicrobial samples due to its agnostic nature and data versatility. However, caveats include high-cost, sequencing depth/sensitivity trade-offs, and challenging bioinformatic deconvolution. Targeted PCR-based profiling optimises sensitivity and cost-effectiveness, but can only identify prespecified targets and may introduce amplification biases. Ultra-high multiplexed PCR is a potential compromise. As no comprehensive comparative evaluation exists, we evaluated performance of each method in taxonomic/resistome-profiling of a well-defined DNA mock sample and seven “real-world” wastewater samples.

**Results:** We tested three sequencing approaches (short-read shotgun metagenomics, Illumina Ampliseq™ ultra-plexed AMR Research Panel, 16S rRNA gene sequencing) with seven bioinformatic pipelines (ResPipe, Illumina DNA Amplicon App, One Codex Metagenomic-/Targeted Loci classification and Ampliseq™ Report, DADA2, and an in-house pipeline for AmpliSeq data [AmpliSeek]). Metagenomics outperformed 16S rRNA gene sequencing in accurately reconstituting a mock taxonomic profile and optimising the identification of diverse wastewater taxa, while 16S rRNA gene sequencing produced more even taxonomic outputs which may be quantitatively inaccurate but enhance detection of low abundance taxa. Shotgun metagenomic and AmpliSeq sequencing performed equally well in profiling AMR genes present in a mock sample, but AmpliSeq identified more genes in more complex, “real-world” samples, likely related to sensitivity of detection at the metagenomic sequencing depth used.

**Conclusions:** A complementary sequencing approach employing 16S rRNA gene or shallow-metagenomic sequencing for taxonomic profiling, and the AmpliSeq AMR panel for high-resolution resistome profiling represents a potential lower-cost alternative to deep shotgun sequencing and may also be more sensitive for the detection of low-prevalence AMR genes. However, our evaluation highlights that both the sequencing and bioinformatics approach used can significantly influence results; for AmpliSeq AMR gene profiling, we developed AmpliSeek which outperformed the other pipelines tested and is open source. Sequencing approach and bioinformatic pipeline should be considered in the context of study goals and sample type, with study-specific validation when feasible.

## Background

Shotgun DNA metagenomic characterisation of polymicrobial samples is increasingly used in both clinical and environmental studies, facilitating agnostic sequencing of all DNA present in a sample and enabling flexible comparisons with reference databases to determine sample composition^1^. For example, the RefSeq^2^ and CARD databases^3^ can be used for taxonomic and antimicrobial resistance (AMR) gene (i.e. “resistome”) profiling respectively. This approach has underpinned multiple studies characterising microbiome-disease associations^4^, evaluating community diversity and anthropogenic impacts^5^, and investigating AMR^6,7^. There is also growing interest in the use of shotgun metagenomics to profile wastewater for population-level surveillance of AMR (i.e. wastewater-based epidemiology [WBE])^8,9^. However, shotgun metagenomics can be expensive, and bioinformatic deconvolution of the data challenging, especially when using short-read sequencing and trying to characterise the genetic context of specific loci, such as AMR genes^10^. Sequencing depth impacts sensitivity to detect rare targets of interest, and therefore, most studies involve a sequencing depth-sensitivity trade-off where uncommon sequences may lack coverage or be missed completely^10^. Additionally, for many studies, only a fraction of the metagenome (e.g. bacterial sequences) will be of interest, and in this context, much of the wider metagenome represents wasted sequencing effort; for example human DNA in studies analysing clinical samples for pathogen diagnostics^1^.

Amplicon-based approaches enrich specific nucleic acid templates during library preparation to increase sensitivity for relevant but less abundant target sequences at reduced cost. A commonly used single target for profiling bacterial communities is the 16S rRNA gene which possesses both highly conserved regions facilitating the use of universal primers, and hypervariable regions which can be used to discriminate between bacterial taxa^11,12^. Increasingly, targeted ultra-highly multiplexed PCR panels have been developed to enable amplicon-based evaluation of the presence/absence and diversity of specific features of a metagenome associated with a key phenotype, for example AMR gene diversity^13-15^, or of a subset of organisms associated with disease, for example respiratory viruses. One example is Illumina AmpliSeq™ which can be used with both Illumina- and community-curated panels (primer pools). These large panels enable capture of a broader range of targets than universal primers or multiplex PCR, whilst theoretically optimising sensitivity and sequencing resource, and are attractive to researchers as they come with defined laboratory protocols and user-friendly bioinformatic pipelines. However, despite internal validation, Illumina-curated panels have demonstrated mispriming events leading to false positive calls of targets; the community-curated panels have largely not undergone any internal validation. Additionally, targeted methods may be prone to amplification biases, and by their nature only focus on specific features of the wider metagenome. Differences in reference database curation and mapping approaches between available bioinformatic pipelines may also influence results^16,17^.

Combining targeted approaches such as 16S rRNA gene sequencing and ultra-highly multiplexed PCR/amplicon sequencing of AMR gene targets represents a potential alternative to deep shotgun metagenomics for profiling species and AMR gene diversity in polymicrobial samples. We therefore assessed the performance of shotgun metagenomics, versus 16S rRNA gene sequencing and the AmpliSeq for Illumina Antimicrobial Resistance Research Panel (henceforth “AmpliSeq”) in reconstituting the true taxonomic composition and resistome of a well-defined DNA mock microbial community. We also compared several bioinformatics approaches to characterising species and AMR gene profiles, including an in-house approach to AmpliSeq data analysis (AmpliSeek). With performance on the mock community as a reference, we then applied the best approach to seven untreated wastewater samples to quantify performance in the context of “real-world” sample complexity. We aimed to highlight potential limitations of each method and recommend potential use cases.

## Methods

### DNA mock community preparation and wastewater samples

Metagenomic, 16S rRNA gene and AmpliSeq sequencing were all conducted on aliquots from the same mock DNA sample (**Fig.1**). The mock was prepared by combining the ZymoBIOMICS Microbial Community DNA Standard (Zymo Research Corporation, Irvine, USA) with three bacterial isolate DNA extracts chosen to enrich the mock sample for clinically relevant AMR genes (**Table S1**); these bacteria had been characterised with whole genome sequencing as part of a previous study evaluating human bloodstream infections^18^.

**Figure 1:**
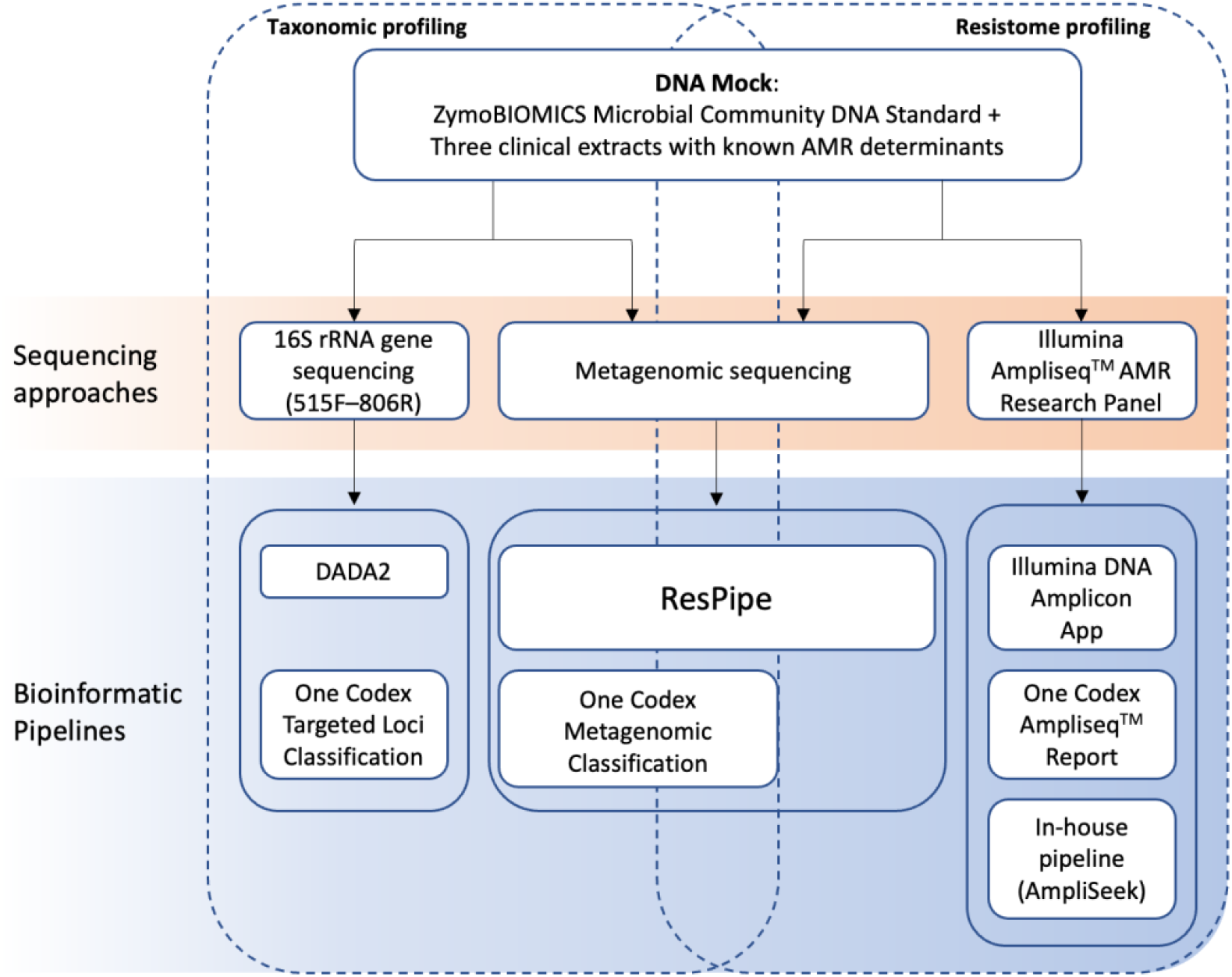
Schematic overview of evaluated sequencing approaches and respective bioinformatic pipelines tested for taxonomic and resistome profiling of the DNA mock.

Wastewater samples comprised seven metagenomic DNA extracts from wastewater influent collected as part of local surveillance in Oxfordshire, UK, 2019, and stored as pellets at -80°C. Metagenomic DNA was extracted using the PowerSoil kit (QIAGEN, Hilden, Germany) according to manufacturer protocols.

### Sequencing

Shotgun metagenomic sequencing on mock DNA community was conducted by Novogene Co (Beijing, China) on the NovaSeq6000 (Illumina, San Diego, USA), generating 150 bp paired-end (PE) reads with a target sequencing depth of 20 million reads (6Gb). AmpliSeq was conducted using the community-designed Illumina for AmpliSeq AMR Research Panel and the Library PLUS kit with manufacturer guidelines; libraries were pooled and sequenced on the Illumina MiniSeq (150 bp PE reads) with replicate libraries prepared for the mock sample. 16S rRNA gene sequencing utilised 515F–806R primers for library preparation and libraries were sequenced on the Illumina MiSeq (250 bp PE reads) as previously described^19^.

Wastewater samples underwent sequencing as described above, however, shotgun sequencing was conducted by the Wellcome Trust Centre for Human Genetics (Oxford, UK) with a depth of ∼75 million reads (∼23Gb) per sample based on previous deep sequencing and rarefaction analyses demonstrating this was the minimum sequencing depth required to capture most AMR gene diversity in this sample type from the same sampling site^10^.

### Software

We tested seven bioinformatic pipelines (**Fig.1, Table S2**). Two pipelines were compared for metagenomic-based taxonomic profiling (ResPipe v1.4.0 and One Codex Metagenomic classification v8/13/2021), two for 16S rRNA-based taxonomic profiling (DADA2 v1.16 [assignTaxonomy] and One Codex Targeted Loci classification v4/15/2021), and three for AmpliSeq-based resistome profiling (Illumina BaseSpace DNA Amplicon App v0.7.12, One Codex AmpliSeq Report v1/17/2019 and an in-house BBTools wrapper pipeline [AmpliSeek] developed as part of this study available at https://github.com/KaibondChau/ampliseek). AmpliSeek performs trimming (BBDuk2) and merging (BBMerge) of reads produced by AmpliSeq sequencing before stringent mapping to AmpliSeq panel target sequences (BBMapSkimmer). Reference databases used by each pipeline are detailed in (**Table S2**).

Geneious Prime v2021.2.2 was used for detailed *in silico* characterisation of the mock AMR profile by visualising the relevant AMR reference sequences and reads mapping to these sequences for each methodological approach (**Fig.S1**).

### Taxonomic profile analysis

We used two previously described error metrics^20^ to quantify differences between theoretical and pipeline-reported distribution of bacterial genera present in the mock sample, a modified mean absolute proportion error (MAPE) based on theoretical vs reported read differences, and Bray-Curtis dissimilarity^21^ (BC) which further considers the total richness present. Both MAPE and BC scores range between 0 and 1, where 0 represents identical composition. For ease of interpretation, scores are presented as 1-MAPE and 1-BC, where higher values represent better accuracy.

Since no theoretical truth existed when characterising wastewater, diversity metrics were used to compare differences between sequencing-pipeline combinations. Chao1 was used as an estimate of taxonomic richness by representing total count of unique observed taxa. Pielou’s evenness was used to assess how similarly represented observed taxa were in abundance estimates (constrained to [0,1] with low values representing unbalanced community estimates where few taxa constitute the majority of abundance). Shannon index was used to quantify both richness and evenness as a measure of overall community complexity where higher values represent increased entropy and therefore complexity. A two-tailed Welch paired *t* test (allowing for different variances in the two samples) was used to determine significance between differences in diversity metrics.

### Sensitivity and specificity calculations for AMR gene detection

The mock DNA community was annotated with AMR target sequences present in the metagenomic and AmpliSeq reference databases using 100% identity threshold to determine a “strict” *in silico* truth. These annotations were compared against pipeline calls using scoring thresholds defined by each pipeline (i.e. thresholds at which AMR targets are called present). For AmpliSeek, scoring thresholds were optimised to maximise sensitivity and specificity based on the “strict” *in silico* truth using receiver operating characteristic curves and Youden’s index^22^ (**Fig.S2**). Established stringent scoring thresholds for AMR target presence were used for ResPipe (lateral coverage=1)^10^ and the One Codex AmpliSeq Report (coverage ≥ 85% and identity ≥ 95%)^23^.

Sensitivity, specificity, positive predictive value (PPV) and negative predictive value (NPV) for AMR target detection were calculated by categorising AMR genes called by each method as true positive, true negative, false positive and false negative, when compared with the *in silico* truth (see above). By adapting this *in silico* truth with respect to the reference AMR gene database used for each method, we fairly compared overall methodological performance (i.e. the CARD v3.0.3 database of 2605 sequences for metagenomic AMR gene profiling approaches and the AmpliSeq reference panel of 815 [AmpliSeek] or 814 [One Codex] amplicon sequences which target 478 AMR genes). However, AmpliSeek scoring thresholds were optimised to the mock dataset and therefore AmpliSeek performance is not directly comparable to the DNA amplicon app and One Codex Ampliseq Report where scoring thresholds were not optimised to the mock dataset.

Mock DNA sequences were also annotated at 98% and 90% identity thresholds to determine “intermediate” and “relaxed” *in silico* truth respectively to investigate AmpliSeq false positives, i.e. to include more AMR gene matches in the truth set as present where these were similar to a gene in the reference database.

## Results

### Taxonomic profiling

All bacterial genera present in the mock DNA community were correctly detected by each sequencing method-pipeline combination, but the relative abundance of different genera was variably under- and overestimated for each combination (**Fig.2A**). This was most notable for 16S rRNA gene-based classification of *Klebsiella* which was underestimated by One Codex Targeted Loci classifier and overestimated by DADA2. *Bacillus* and *Lactobacillus* were also inaccurately estimated for One Codex Targeted Loci and Metagenomic estimates respectively. Regardless of pipeline, shotgun metagenomics more accurately reconstituted true taxonomic composition over 16S rRNA gene-sequencing (**Fig.2B**; mean 1-MAPE=0.77 vs 0.54, and mean 1-BC=0.87 vs 0.75). Differences in performance between pipelines within each dataset was small, with ResPipe (1-MAPE:0.79; 1- BC:0.88) marginally outperforming One Codex metagenomic classifier (0.75; 0.86) for shotgun sequencing data, and DADA2 (0.55; 0.77) outperforming the One Codex Targeted Loci classifier 0.52; 0.73) for 16S rRNA gene sequencing data (**Fig.2B**).

**Figure 2:**
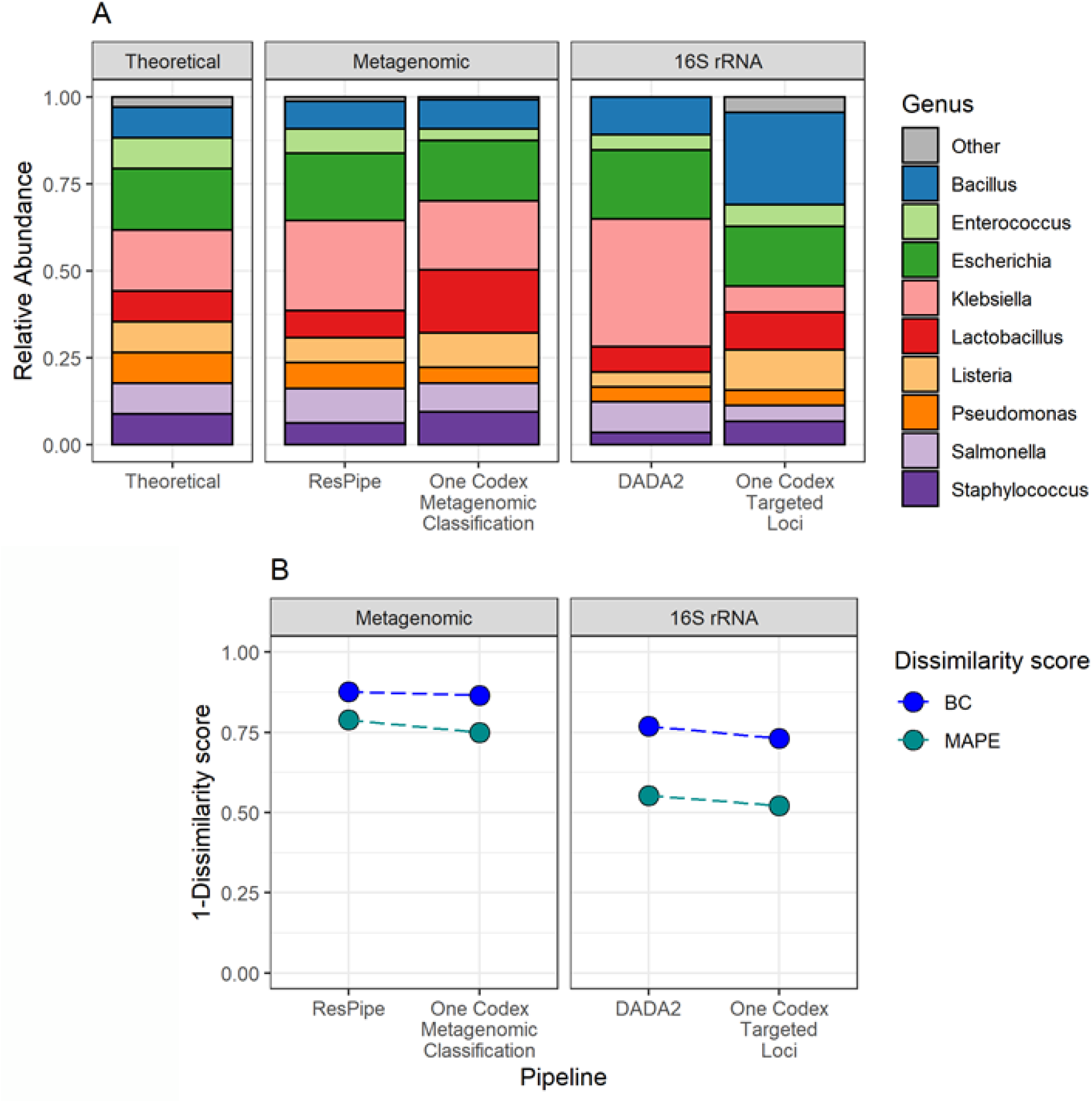
(A) Theoretical taxonomic composition of DNA mock as compared to actual generated pipeline outputs faceted by sequencing approach. Genus “Other” represents genera present at <1% relative abundance in pipeline outputs. For the theoretical facet, “Other” represents DNA composition from yeasts included in the commercial DNA standard. (B) Taxonomic profiling error scores represented by 1-BC and 1-MAPE where 1.00 is perfect reconstitution of theoretical genera abundances.

### AMR gene profiling

Annotation of the mock DNA sequences with approach-specific reference databases identified 77 CARD and 43 AmpliSeq target sequences present at 100% similarity. These were all detected with 100% sensitivity by all respective methods/pipelines (**Fig.3A**), except the DNA Amplicon App, which performed poorly (22/43 known AmpliSeq targets detected; 51%). For precision, ResPipe (specificity=0.99; PPV=0.72) performed best, followed by AmpliSeek (0.98; 0.69), DNA Amplicon App (0.98; 0.65) and One Codex AmpliSeq Report (0.94; 0.49) (**Fig.3B**, **3C**). However, notably, we found that most false positive hits obtained (67/75; 89%) arose from ≤2% nucleotide variation from reference AMR sequences (AmpliSeek [18/19], DNA Amplicon App [12/12] and One Codex AmpliSeq Report [37/44]) (**Fig.4**).

Replicate AmpliSeq libraries produced consistent results with no impact on performance metrics for all pipelines tested (**Fig.S3**).

AmpliSeq false positives were investigated by comparing pipeline calls to “intermediate” and “relaxed” mock *in silico* truth (see Methods). We considered AmpliSeq reference sequences present in the mock with 98-100% identity as target variants (**Fig.4**) and assessed these separately to account for the limited sequence diversity amongst reference sequences for specific targets. An additional 37 reference sequences were identified in the mock at 98-100% identity. When these target variants were included in overall performance metrics, One Codex AmpliSeq Report retained 100% sensitivity with increased specificity and PPV (0.99; 0.92). However, AmpliSeek and DNA Amplicon App were less able to detect target variants, with overall sensitivities of 0.76 and 0.43 respectively. However, AmpliSeek sensitivity improved to 0.90 (**Fig.4** - grey fill) when including less stringent scoring thresholds (i.e. “potentially detected” calls) (**Fig.S1**).

**Figure 3:**
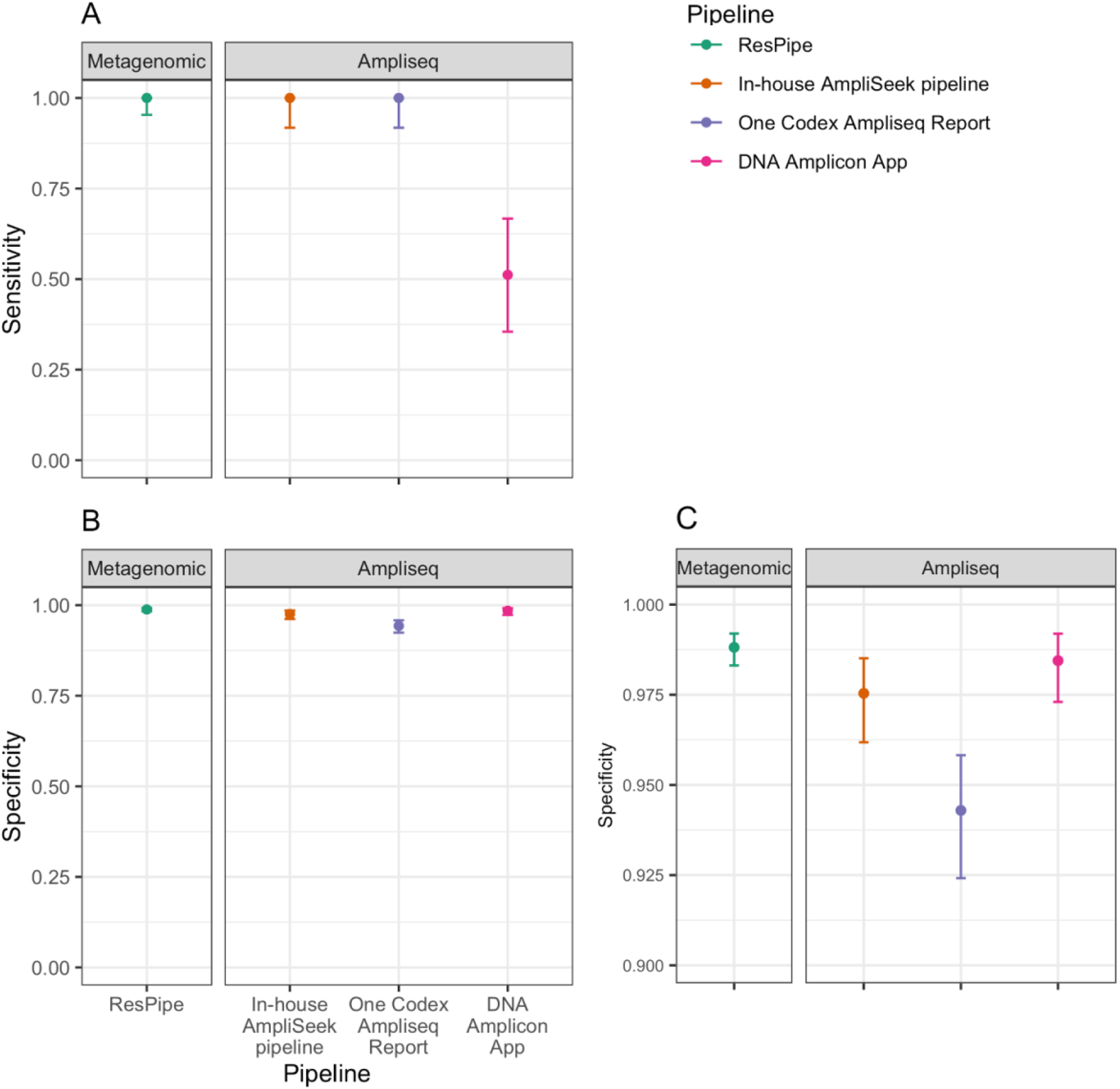
(A) Detection sensitivity of known AMR genes present in the mock at 100% identity. NB the reference AMR gene database used to annotate the mock was specific to, and hence different for, each approach. Error bars represent 95% confidence intervals (CIs), and facets divide sequencing approach and bioinformatic pipelines. **(B) Specificity of AMR gene detection presented as for (A) and with magnified y-axis in (C)**.

**Figure 4:**
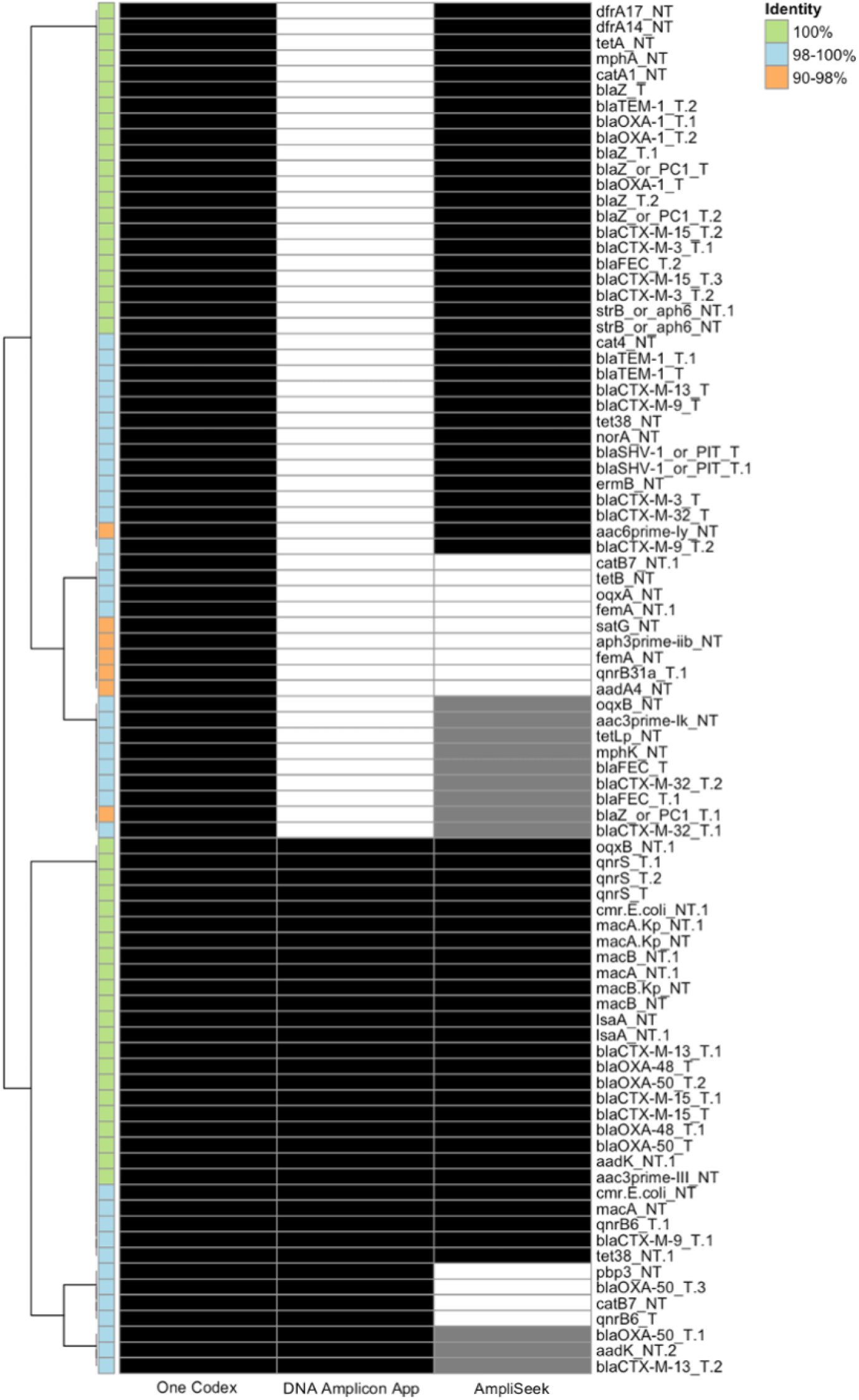
Heatmap of AmpliSeq pipeline calls labelled by reference marker ID and sequence identity. Each column represents a bioinformatic pipeline output where black and grey fill reflect detected and potentially detected calls respectively. Labels (green, blue, orange) denote the sequence identity of reference markers when annotated in the mock DNA community.

### Real-world wastewater samples

#### Taxonomic profiles

Consistent with findings from our mock microbial community benchmarks, taxonomic profiles of “real-world” wastewater samples differed significantly between 16S rRNA and shotgun sequencing, with large differences in relative abundance and detection of core taxa (**Fig.5**). For metagenomics, Proteobacteria were estimated as most abundant across all samples, with relative abundance approximately two-fold higher than reported by 16S rRNA gene sequencing. Conversely, relative abundance of phyla detected by 16S rRNA gene sequencing across all samples were reduced in metagenomic estimates (Firmicutes, Bacteriodota) or entirely missed/found at <1% relative abundance (Fusobacteriota, Campylobacteriota). Interestingly, this was not the case for Actinobacteriota which was estimated with higher relative abundance than 16S rRNA gene-based estimates by both shotgun pipelines. 16S rRNA gene-based estimates of phylum abundance were more even, with variable detection of Chloroflexi, Patescibacteria, Planctomycetota and Verrucomicrobiota across samples. Interestingly, Campylobacteriota was not detected using the One Codex Targeted Loci pipeline but reported by DADA2. Alternative faceting to highlight difference on a per-sample basis is provided in (**Fig.S4**).

**Figure 5:**
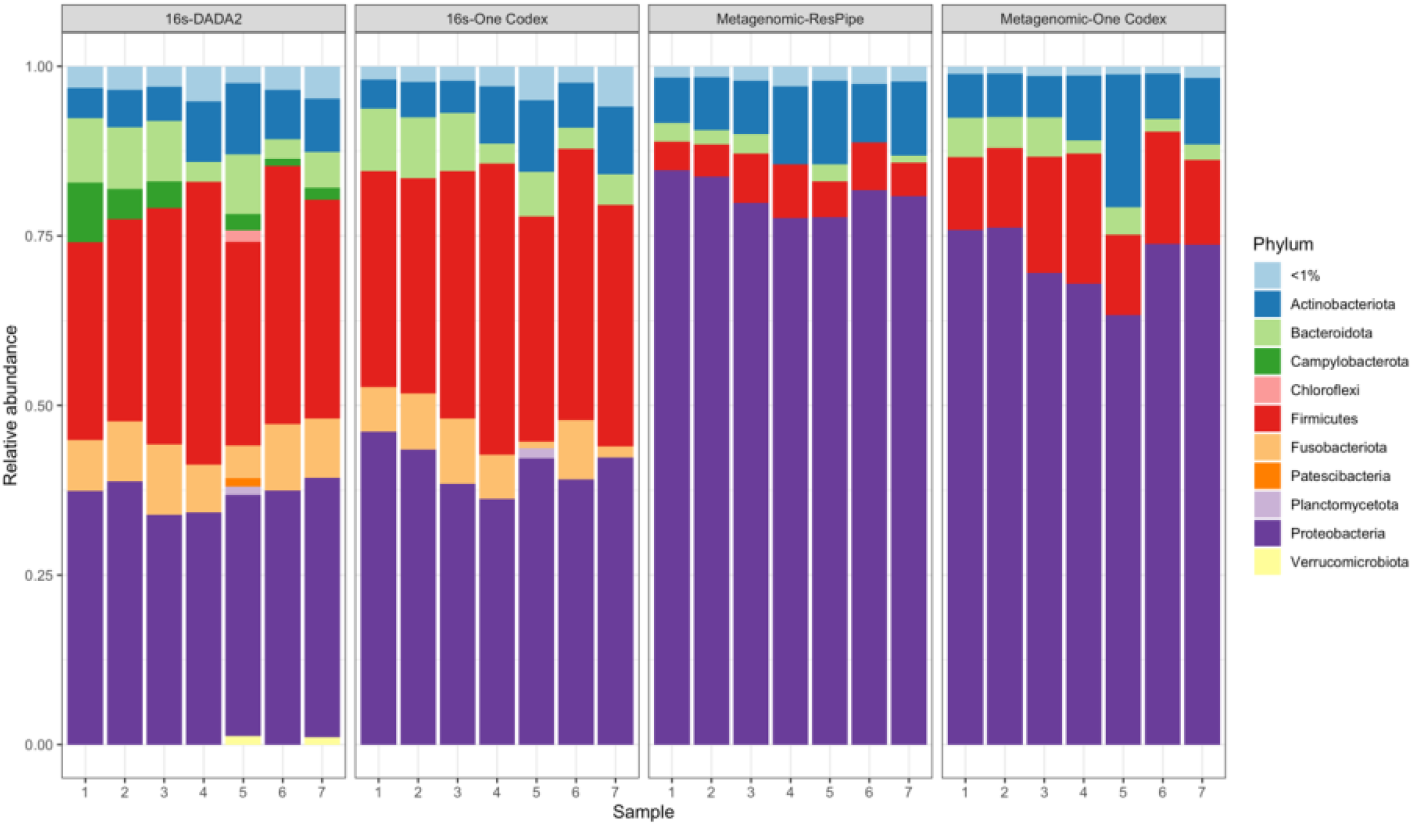
Relative abundance of phyla for wastewater samples (1-7) faceted by sequencing approach and pipeline combination.

Metagenomic approaches consistently reported significantly higher richness (Chao1) than 16S rRNA gene sequencing approaches at all taxonomic ranks tested (p<0.002; mean increase of 1354 genera), with the One Codex Metagenomic Classifier identifying significantly more unique taxa than ResPipe (p<0.001; mean increase of 787 genera) (**Fig.6A**). Conversely, 16S rRNA gene sequencing approaches produced profiles with significantly more even representation of taxa (Pielou’s evenness; p<0.001) than metagenomic approaches (**Fig.6B**). Despite higher richness in metagenomic results, overall community complexity (Shannon index) was significantly higher for profiles generated by 16S rRNA gene sequencing than metagenomic approaches (p<0.007); except for ResPipe at the genus level. (**Fig.6C**). Interestingly, One Codex Targeted Loci classifier (16S rRNA gene data) produced taxonomic profiles with higher variability across the wastewater samples than all other sequencing-pipeline combinations (**Fig.6ABC**).

**Figure 6:**
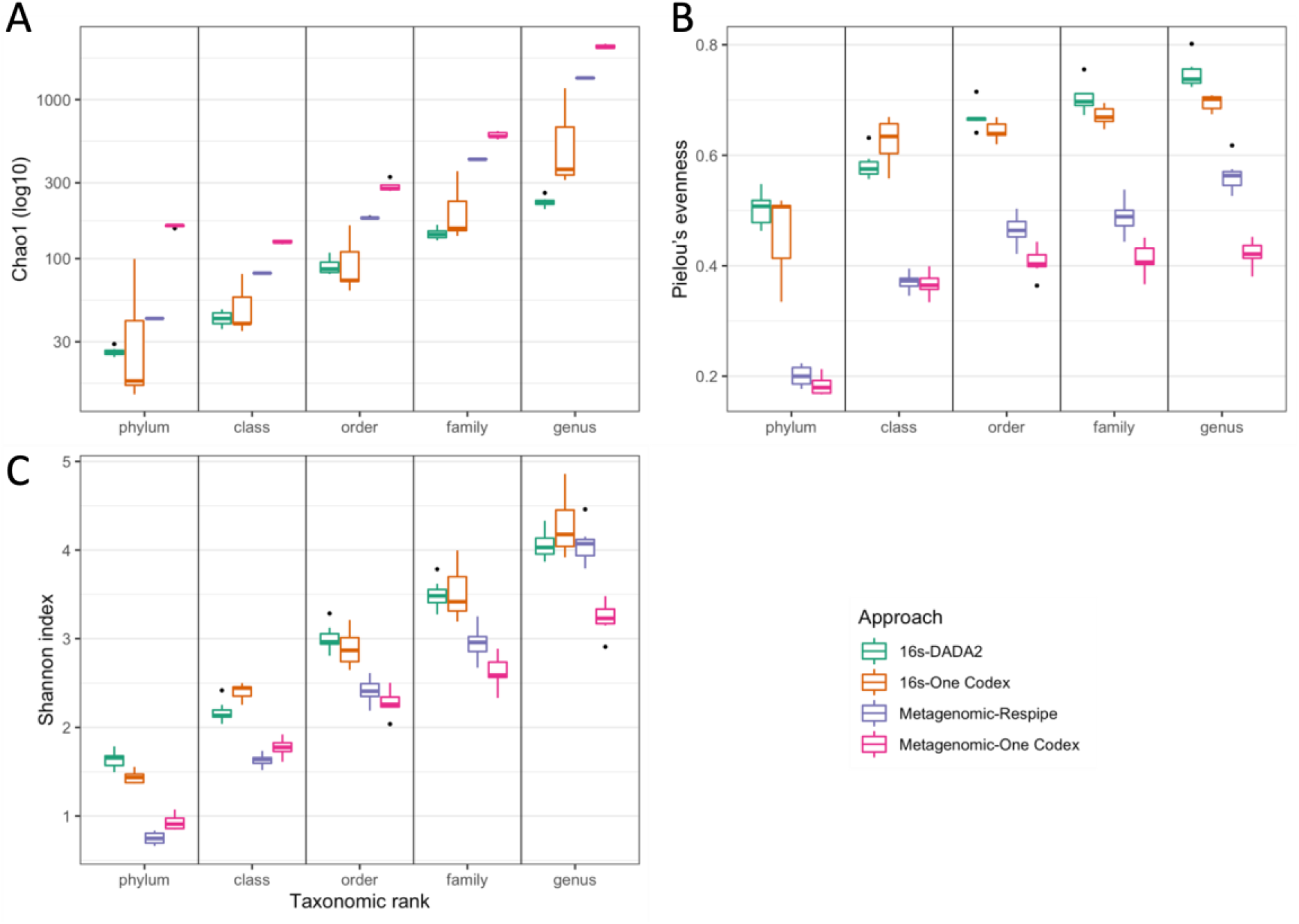
Distribution of diversity metrics across all wastewater samples coloured by sequencing approach and pipeline combination. Panels: 6A reports richness via log10 Chao1, 6B reports abundance distribution via Pielou’s evenness, 6C reports overall alpha diversity via Shannon index.

#### AMR profiles

Across all wastewater samples, AmpliSeq One Codex Report identified a total of 3220 AmpliSeq targets (2889 called as present, 331 called as probable) while AmpliSeek identified a total of 2157 (1970 called as present, 187 called as potentially present), and the shotgun metagenomics-ResPipe pipeline identified a total of 526 CARD sequences with lateral coverage of 1 (i.e. 100% nucleotide identity over the full length of the reference sequence). When deduplicated to account for targets shared across wastewater samples and redundant AmpliSeq amplicon design (**Fig.S1**), the AmpliSeq One Codex Report detected the most unique AMR genes (n=367), followed by AmpliSeek (n=300) and shotgun metagenomics-ResPipe (n=132), respectively.

The distributions of AmpliSeq targets identified by the One Codex Report and AmpliSeek showed the same ratio of increased present calls as the evaluation of the defined mock (i.e. 1.4-fold more AmpliSeq targets called as present by One Codex than AmpliSeek). Similarly consistent with the methodological comparisons on the mock sample, almost all AMR genes called as present by One Codex but absent by AmpliSeek were hits identified at <100% nucleotide identity (817/826; 99%; median nucleotide identity: 98% [IQR:97-99%]). The nine targets identified as present and at 100% identity by the One Codex pipeline but called absent by AmpliSeek possessed low numbers of mapped reads by One Codex (median of one read mapping only). The single AmpliSeq target not targeted by the One Codex AmpliSeq Report (aph2prime-Ia_NT) was called as present by AmpliSeek across all wastewater samples.

## Discussion

In this study, we have compared the performance of metagenomic and targeted sequencing and different bioinformatics approaches in accurately reconstructing taxonomic and AMR gene composition of a defined mock microbial community, and a set of polymicrobial “real-world” wastewater samples containing complex microbial communities. We highlight that results are significantly influenced by both the sequencing and bioinformatics approach used, and this is an important consideration for studies using these methods.

### Mock community evaluation

Metagenomics outperformed targeted 16S rRNA sequencing and more accurately estimated true relative abundance of mock genera regardless of the bioinformatic pipeline utilised. While 16S rRNA gene sequencing successfully captured all mock genera, relative abundances differed significantly from the known, true, composition. This was not unexpected as targeted approaches are subject to primer and amplification biases already known to potentially distort taxonomic profiles^24-26^, however, the degree of bias has not been previously quantified. This was most apparent for One Codex Targeted Loci results where *Klebsiella* and *Bacillus* abundances were respectively under- and overestimated by up to three-fold. Conversely, shotgun metagenomics is less susceptible to these biases, and when combined with statistical methods for normalising abundances (as used by both ResPipe and One Codex Metagenomic Classifier), accurately estimated mock genus distributions. For profiling taxonomy, ResPipe and DADA2 slightly outperformed the One Codex Metagenomic and Targeted Loci classifiers. Since classification methods were similar within each sequencing approach (i.e. k-mer based for shotgun metagenomics and alignment-based for 16S rRNA gene profiling), differences in the reference taxonomic databases are most likely responsible for differences in performance^16^, consistent with previous studies demonstrating classification is dependent on database choice^20,24^.

Almost all metagenomics and AmpliSeq approaches detected all relevant unique AMR genes known to be present in the mock microbial community. However, the DNA Amplicon App performed poorly, reporting no reads mapping to 21 AmpliSeq target sequences known to be present in the mock and reported as present by the One Codex AmpliSeq Report and AmpliSeek. This appears attributable to differences in the read mapping workflow rather than alignment algorithm *per se* since both One Codex AmpliSeq Report and the DNA Amplicon App use the BWA algorithm to align reads. However, instead of directly aligning reads to the reference AmpliSeq target sequences as performed by One Codex and AmpliSeek, the DNA Amplicon App first aligns reads to a reference database of whole genomes containing the reference AMR genes^27^. We noted that this initial genome-based screen appeared to erroneously filter out reads which would otherwise map directly to sequences present in the AmpliSeq AMR panel. However, full details of the DNA Amplicon App workflow are unclear and detailed evaluation of this pipeline was therefore beyond the scope of this study.

For AmpliSeq based methods, when focussing on AMR genes known to be present in the mock with 100% nucleotide sequence identity, AmpliSeek slightly outperformed One Codex AmpliSeq Report as the latter reported more false positives. False positives have been previously described with the AmpliSeq panels and attributed to mispriming events associated with primers erroneously binding to non-target sequences^28^. However, mispriming did not appear to be the main cause in this study since most false positives shared high sequence identity to the reference AMR genes (>98%), indicating that these are more likely related to pipeline-specific differences in calling genes present/absent. In fact, the highly related nature of most false positive calls made by the One Codex AmpliSeq Report would be consistent with the default 2% error BWA mapping threshold used in the pipeline to account for sequencing error. The difficulty in calling presence/absence of related AMR genes where the reference sequences used as markers are very similar is highlighted in our analyses: For example, the amplicon reference sequence for *bla*_FEC_ (blaFEC_T) shares high nucleotide identity with amplicon reference sequences denoting *bla*_CTX-M-3_, *bla*_CTX-M-15_ and *bla*_CTX-M-32_ (blaCTX-M-3_T.2 [99%], blaCTX-M- 15_T.3 [99%] and blaCTX-M-32_T.1 [98%]). Modifying the mapping threshold may avoid these issues since recent work has suggested the median error rate of Illumina MiniSeq sequencing reads may be as low as 0.6%^29^. For AmpliSeek we retained stringent mapping of reads as the default to enable differentiation between highly similar reference markers, guard against mispriming, and call specific alleles with high confidence, but this threshold can be relaxed depending on the users’ requirements.

#### “Real-world” wastewater samples

On real-world samples, where the true diversity was much higher than in the mock community, differences in the relative abundance of taxa and the number of unique AMR genes observed between approaches when profiling the mock community were greatly amplified. In these samples, metagenomic approaches identified more unique taxa than 16S rRNA gene profiling at all taxonomic ranks tested. This represents a commonly cited key advantage of shotgun sequencing, where higher richness of bacterial taxa can be reported than for 16S rRNA-based methods^30-32^, and is particularly important in the characterisation of complex microbial communities^31^. Taxonomic classification is also heavily dependent on the reference database used^17^. In this respect, the One Codex Metagenomic Classifier yielded significantly higher richness than ResPipe likely owing to respective reference species database sizes of ∼115,000 and ∼20,000 complete genomes. Taxonomic richness reported by ResPipe was also seemingly capped across most ranks with minimal variation between wastewater samples which may be a result of saturating the reference database as previously suggested^10^.

Metagenomic abundance estimates were largely skewed towards a small number of taxa; in contrast, 16S rRNA gene-based estimates were consistently more even across all ranks regardless of pipeline, and more sensitive to the detection of taxa missed by shotgun metagenomic outputs. This likely partly reflects the sensitivity trade-off of shotgun sequencing where sequencing effort can be swamped by high abundance taxa and/or taxa with large genomes; especially when sequencing depth is low and community complexity is high^1^, which can result in missing the presence of low-abundance, but significant, taxa^33^. However, users of 16S rRNA gene sequencing for taxonomic profiling need to be aware of the likely greater divergence of profiles from the truth. In our hands, different 16S rRNA gene pipelines also notably impacted results; for example, Campylobacteriota were absent from One Codex outputs but classified as present using DADA2, further highlighting differences across reference databases.

Interestingly, despite being more sensitive in profiling AMR targets in the mock, metagenomics detected less than half the number of AMR genes reported by targeted AmpliSeq approaches. This is potentially driven by low abundance AMR genes which are missed by an agnostic approach, but directly targeted and amplified by AmpliSeq. However, we used the strictest definition of AMR gene presence requiring read matches covering all of the reference sequence (lateral coverage=1), so our metagenomic profiling was a conservative estimate. As discussed, sequencing depth impacts the sensitivity of target detection in complex samples, with the relative simplicity of the mock composition (n=11 bacterial genomes, 77 CARD sequences, 43 AmpliSeq sequences) masking this limitation.

As seen in the evaluation on the mock, the less stringent mapping thresholds of the One Codex Report produced more present calls than AmpliSeek due to calling target variants as present. In fact, the proportion of increased present calls made by One Codex Report compared to AmpliSeek was consistent between mock results and wastewater results. These findings support similar performance of both AmpliSeq pipelines in wastewater as in the mock despite significantly increased complexity, where AmpliSeek results represent increased confidence in the presence of the exact marker sequence.

#### Limitations

We have not evaluated all methods for profiling taxonomic and resistome composition and therefore potentially missed approaches with increased performance. However, our pragmatic evaluation focussed on accessible approaches (i.e. kit-based and all-in-one pipelines) which are commonly used, attractive to researchers and have not undergone comparative evaluation. The relative simplicity of the mock DNA community may mask sensitivity issues owing to sequencing depth but use of *in silico* simulated datasets would otherwise omit assessment of library preparation amplification reactions. To mitigate this, we enriched the mock with additional AMR determinants and presented performance differences between the mock and “real-world” samples. Our in-house pipeline AmpliSeek was internally validated on the same mock dataset used for performance metrics which inherently optimises performance, however, the “real-world” samples represented an external validation confirming similar performance with and without scoring threshold optimisation. We also guarded against wastewater metagenomic sensitivity issues by using optimised depth as determined by previous ultradeep pilot sequencing performed on wastewater form the same sampling site^10^. Our resistome characterisation focusses on presence/absence but a major advantage of shotgun metagenomics is quantitative measurement of genes which we have not explored here. Additional factors such as read length and approaches to normalization may also optimise each approach but exploring these were beyond the scope of this study.

## Conclusion

A complementary sequencing approach encompassing 16S rRNA or shallow-metagenomic sequencing and the AmpliSeq AMR panel represents a potential lower-cost alternative to deep shotgun sequencing for taxonomic and resistome profiling of highly diverse, complex, polymicrobial samples. Accuracy of profiling was also to some extent dependent on the bioinformatics pipeline used; for AmpliSeq AMR gene profiling we have optimised an in-house pipeline (AmpliSeek) which outperformed the other methods tested on this dataset and is open source. However, further validation of this in-house pipeline is needed, since it was also developed on the same dataset.

We recommend careful consideration and further validation of sequencing approaches and bioinformatic pipelines in the context of study goals and sample type to be analysed. Validating sequencing and bioinformatic approaches remains challenging due to difficulties in representing the true complexity of “real-world” samples; any conclusions drawn should always be considered in the context of the methods used.

## Funding

This work was supported by the Medical Research Foundation National PhD Training Programme in Antimicrobial Resistance Research, the National Institute for Health Research (NIHR) Health Protection Research Unit in Healthcare Associated Infections and Antimicrobial Resistance at University of Oxford (NIHR200915) in partnership with the UK Health Security Agency (UKHSA), and the NIHR Oxford Biomedical Research Centre. The report presents independent research. The views expressed in this publication are those of the authors and not necessarily those of the NHS, NIHR, UKHSA or the Department of Health and Social Care.

KKC and WM are Medical Research Foundation PhD students (ref. MRF-145-0004-TPG-AVISO). NS is an Oxford Martin School fellow and an NIHR Oxford BRC Senior Research Fellow. DWE is a Robertson Foundation Fellow and an NIHR Oxford BRC Senior Fellow. ASW is an NIHR Senior Investigator.

## Contributors

A complete list of author contributions as per CRediT; Contributor Roles Taxonomy is given in the appendix.

## Availability of data and materials

Raw sequencing data are available under ENA project PRJEB52722 with all metadata included in supplementary file 1. R scripts and markdown renders are available at https://github.com/KaibondChau/Chau_etal_Ampliseek. The AmpliSeek pipeline is available at https://github.com/KaibondChau/ampliseek.

## Notes

### Competing Interest Statement

The authors have declared no competing interest.

