## appendix for "Evaluation of metagenomic, 16S rRNA gene and ultra-plexed PCR-based sequencing approaches for profiling antimicrobial resistance gene and bacterial taxonomic composition of polymicrobial samples"

**Table S1: Mock DNA community theoretical composition**

| **Species (source)** | **Theoretical genomic DNA composition (%)** | **ResFinder WGS-predicted resistance** |
| --- | --- | --- |
| *Pseudomonas aeruginosa* (ZymoBIOMICS^TM^) | 8.82 | aph(3')-IIb (aph(3')-IIb_CP006832); blaPAO (blaPAO_AY083592); blaPAO (blaPAO_AY083595); catB7 (catB7_AF036933); crpP (crpP_HM560971) |
| *Escherichia coli* (ZymoBIOMICS^TM^) | 8.82 | sitABCD (sitABCD_AY598030) |
| *Salmonella enterica* (ZymoBIOMICS^TM^) | 8.82 | aac(6')-Iaa (aac(6')-Iaa_NC_003197) |
| *Lactobacillus fermentum* (ZymoBIOMICS^TM^) | 8.82 | None |
| *Enterococcus faecalis* (ZymoBIOMICS^TM^) | 8.82 | lsa(A) (lsa(A)_AY225127) |
| *Staphylococcus aureus* (ZymoBIOMICS^TM^) | 8.82 | aadD (aadD_M19465); blaZ (blaZ_JBTH01000015); tet(L) (tet(L)_M29725); qacG (qacG_EU622633) |
| *Listeria monocytogenes* (ZymoBIOMICS^TM^) | 8.82 | fosX (fosX_AL591981) |
| *Bacillus subtilis* (ZymoBIOMICS^TM^) | 8.82 | None |
| *Saccharomyces cerevisiae* (ZymoBIOMICS^TM^) | 1.47 | None |
| *Cryptococcus neoformans* (ZymoBIOMICS^TM^) | 1.47 | None |
| *Escherichia coli* (clinical isolate)  ENA: ERS11963948 | 8.82 | aac(3)-IId (aac(3)-IId_EU022314); aadA5 (aadA5_AF137361); aph(3'')-Ib (aph(3'')-Ib_AF321551); aph(6)-Id (aph(6)-Id_M28829); aadA5 (aadA5_AF137361); blaTEM-1B (blaTEM-1B_AY458016); blaCTX-M-14 (blaCTX-M-14_AF252622); catA1 (catA1_V00622); dfrA17 (dfrA17_FJ460238); erm(B) (erm(B)_JN899585); mph(A) (mph(A)_D16251); mcr-5.1 (mcr-5.1_KY807921); mph(A) (mph(A)_D16251); qacE (qacE_X68232); sitABCD (sitABCD_AY598030); sul2 (sul2_HQ840942); sul1 (sul1_U12338); tet(B) (tet(B)_AF326777) |
| *Klebsiella pneumoniae* (clinical isolate)  ENA: ERS11963949 | 8.82 | aadA2 (aadA2_JQ364967); aph(3'')-Ib (aph(3'')-Ib_AF024602); aph(3'')-Ib (aph(3'')-Ib_AF313472); aph(3'')-Ib (aph(3'')-Ib_AF321550); aph(3'')-Ib (aph(3'')-Ib_AF321551); aph(6)-Id (aph(6)-Id_M28829); blaLAP-2 (blaLAP-2_EU159120); blaOXA-48 (blaOXA-48_AY236073); blaSHV-145 (blaSHV-145_JX013655); blaSHV-179 (blaSHV-179_KF705208); blaSHV-194 (blaSHV-194_KX421191); blaSHV-199 (blaSHV-199_MF373391); blaSHV-26 (blaSHV-26_AF227204); blaSHV-78 (blaSHV-78_AM176553); blaSHV-98 (blaSHV-98_AM941844); dfrA12 (dfrA12_AM040708); fosA (fosA_AFBO01000747); OqxA (OqxA_EU370913); OqxB (OqxB_EU370913); qacE (qacE_X68232); qnrS1 (qnrS1_AB187515); sul1 (sul1_U12338); sul2 (sul2_AY034138); tet(A) (tet(A)_AJ517790) |
| *Klebsiella pneumoniae* (clinical isolate)  ENA: ERS11963950 | 8.82 | aac(3)-IIa (aac(3)-IIa_CP023555) ; aac(6')-Ib-cr (aac(6')-Ib-cr_DQ303918); aac(6')-Ib-cr (aac(6')-Ib-cr_DQ303918) ; aph(3'')-Ib (aph(3'')-Ib_AF321551); aph(6)-Id (aph(6)-Id_M28829) ; blaCTX-M-15 (blaCTX-M-15_AY044436); blaCTX-M-15 (blaCTX-M-15_AY044436) ; blaOXA-1 (blaOXA-1_HQ170510); blaOXA-1 (blaOXA-1_HQ170510) ; blaSHV-164 (blaSHV-164_HE981194); blaSHV-164 (blaSHV-164_HE981194) ; blaSHV-59 (blaSHV-59_AY790341) ; blaTEM-1B (blaTEM-1B_AY458016); blaTEM-1B (blaTEM-1B_AY458016) ; catB3 (catB3_AJ009818); catB3 (catB3_U13880) ; dfrA14 (dfrA14_KF921535); fosA (fosA_ACZD01000244) ; OqxA (OqxA_EU370913); OqxB (OqxB_EU370913); OqxB (OqxB_EU370913) ; qnrB1 (qnrB1_DQ351241); sul2 (sul2_AY034138) ; tet(A) (tet(A)_AJ517790) |

**Table S2: Bioinformatic pipeline versions and associated reference databases**

| **Sequencing approach** | **Profiling strategy** | **Bioinformatic pipeline** | **Reference database** |
| --- | --- | --- | --- |
| Shotgun metagenomics | Taxonomy | ResPipe v1.4.0 | RefSeq "Complete genome" assembly level sequences v91 |
|  |  | One Codex Metagenomic classification v8/13/2021 | One Codex Genomes database |
|  | Resistome | ResPipe v1.4.0 | CARD v3.03 |
| 16S rRNA | Taxonomy | One Codex Targeted Loci classification v4/15/2021 | One Codex Targeted Loci database |
|  |  | DADA2 v1.16 | SILVA SSU |
| AmpliSeq | Resistome | Illumina BaseSpace DNA Amplicon App v0.7.12 | Illumina AmpliSeq panel reference genomes and manifest target sequences v1.0 |
|  |  | One Codex AmpliSeq Report v1/17/2019 | One Codex-curated AmpliSeq marker sequences |
|  |  | In-house pipeline (AmpliSeek) | Illumina AmpliSeq panel manifest target sequences v1.0 |

**Fig.S1: Example visualisation of AmpliSeq amplicon target sequences used for generating *in silico* truth**

AmpliSeq target sequences for bla_CTX-M-15_ annotated onto sequences from the mock DNA community at 100% sequence identity using Geneious. NB AmpliSeq panel design is degenerate where multiple primer pairs target different positions on the same gene – denoted by a T.# suffix.

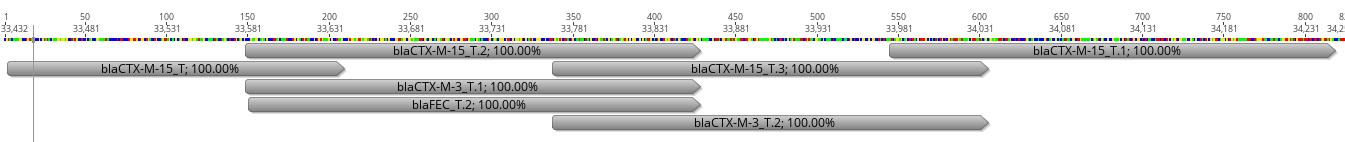

**Figure S2: AmpliSeek scoring thresholds**

Left: Receiver operator characteristic curve for mock AmpliSeek output metrics in relation to true positives and false positives based on the known “strict” *in silico* truth. Top right: Optimal thresholds as determined by Youden’s index. Lnorm_count_prop denotes the proportion of length-normalised count for a specific AmpliSeek target. Bottom right: Combined thresholds to determine target status.

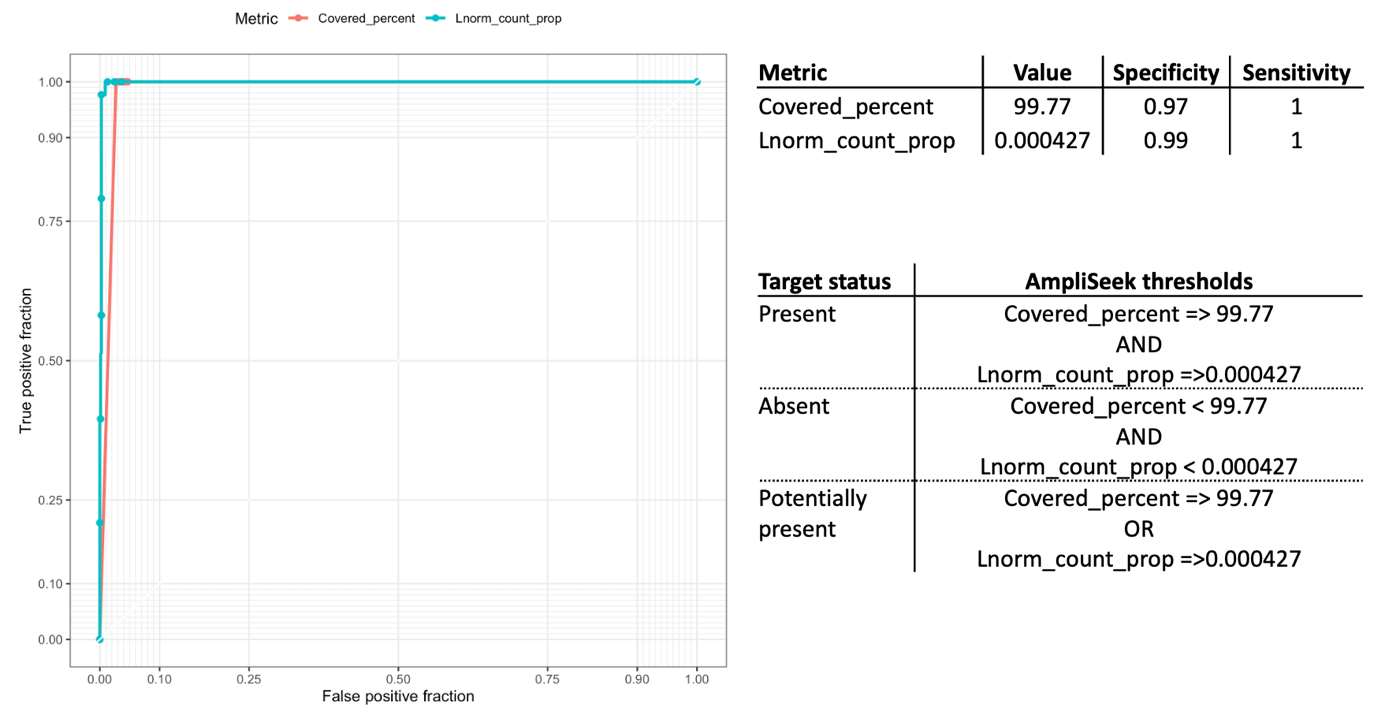

**Figure S3. Replicate results of AmpliSeq profiling.** Two aliquots of mock community DNA were independently prepared for AmpliSeq (see main text methods) to assess replicability of ultra-high multiplexed PCR and library prep. Read counts for AMR targets between replicates were highly concordant for both the DNA Amplicon App (Lin's concordance correlation coefficient [CCC]=0.99) (panel A) and in-house AmpliSeek (CCC=1.00) (panel B). Interestingly, the read difference per AMR target between replicates was reduced in AmpliSeek (median=502; IQR: 14-2162) compared to the DNA Amplicon App (median=1746; IQR: 304-5466). Disagreements (i.e. reads mapping to a target in only one replicate) were rare for both DNA Amplicon App (n=2) and AmpliSeek (n=1), and associated to very low read counts (≤2). Variations in read count or target coverage did not impact overall performance metrics for presence/absence calls for either pipeline. Read-level data was not available from the One Codex AmpliSeq report, but overall performance metrics did not differ for One Codex replicates either.

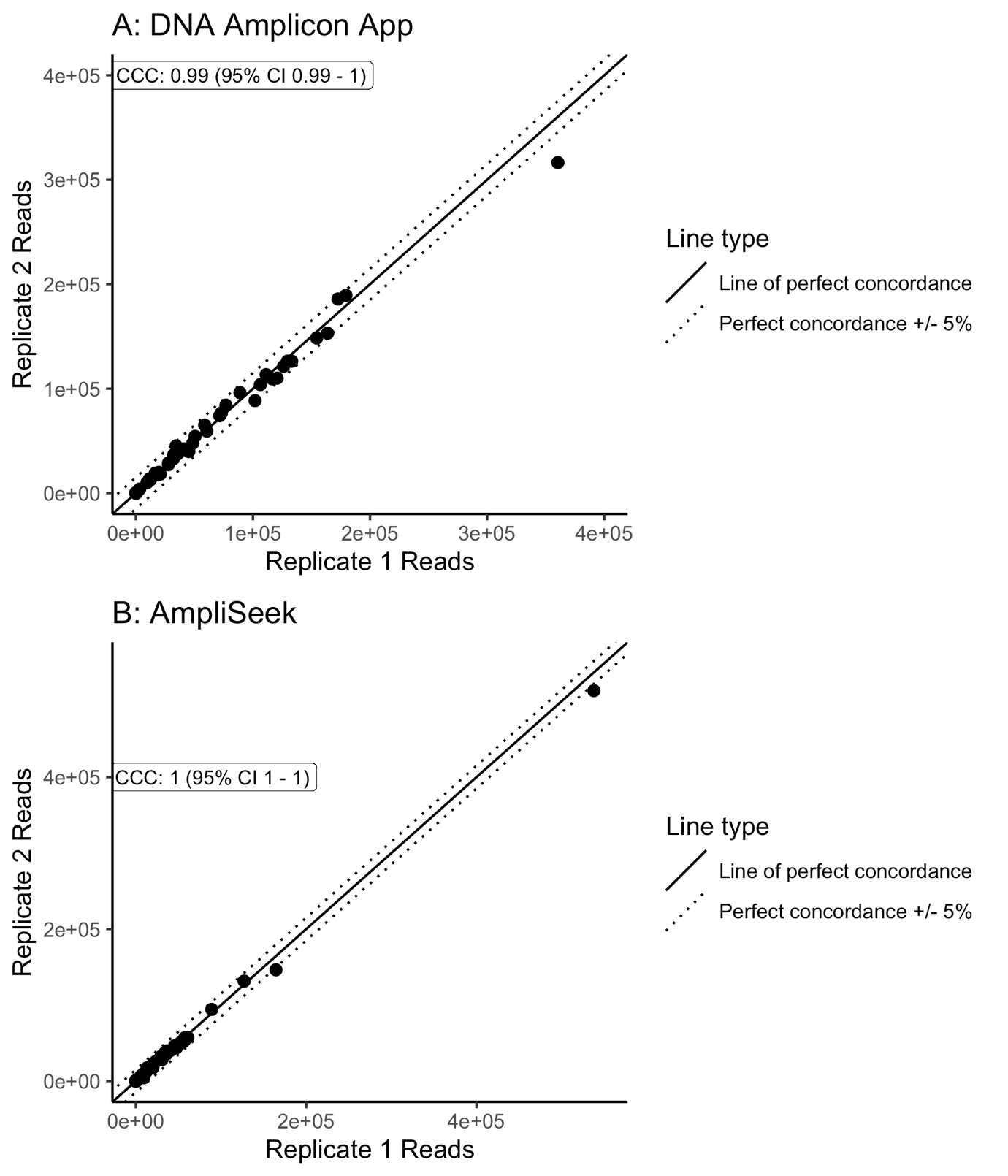

**Figure S4: Relative abundance of phyla for wastewater samples with alternative faceting by wastewater samples.**

Wastewater samples are denoted by 1-7. Sequencing-pipeline approach combinations are denoted as follows: a = 16S-DADA2; b = 16S-One Codex; c = Metagenomic-ResPipe; d=Metagenomic-One Codex.

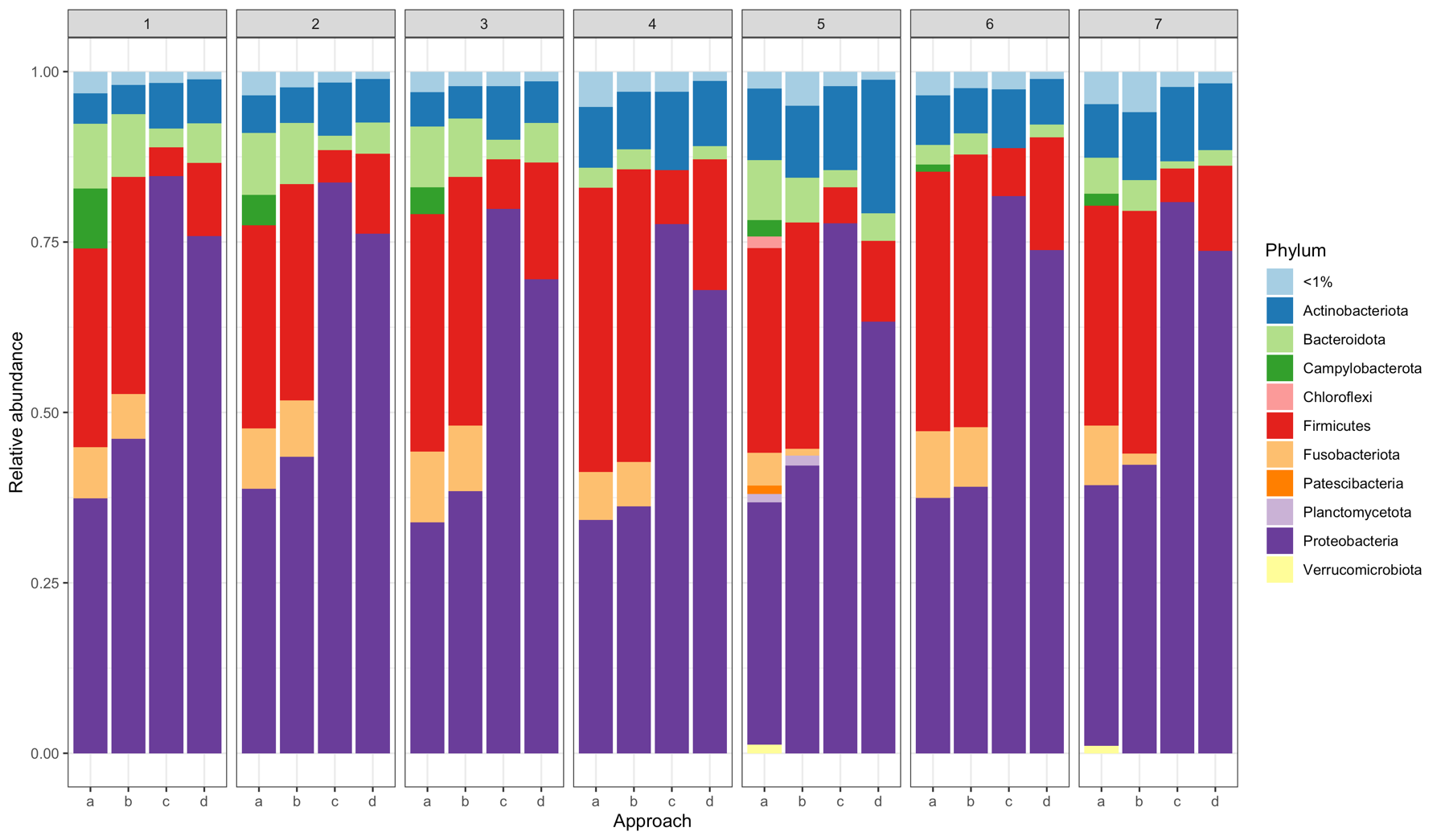

**Complete Author contributions (as per CRediT; Contributor Roles Taxonomy)**

**Conceptualization**

KK Chau, DS Read, N Stoesser, AS Walker

**Methodology**

KK Chau, DS Read, N Stoesser

**Software**

KK Chau, W Matlock, B Constantinides, DW Eyre

**Validation**

KK Chau, N Stoesser

**Formal Analysis**

KK Chau, N Stoesser

**Investigation**

KK Chau, N Stoesser, Newbold L, Tipper H, Goodall T

**Resources**

KK Chau, Lipworth S, Brett H, Hughes J

**Data curation**

KK Chau, W Matlock, B Constantinides, N Stoesser

**Writing – original draft**

KK Chau, N Stoesser

**Writing – review & editing**

KK Chau, N Stoesser, DS Read, AS Walker, DW Eyre

**Visualization**

KK Chau, N Stoesser

**Supervision**

KK Chau, DW Crook, DS Read, N Stoesser, AS Walker

**Project administration**

KK Chau, DS Read, N Stoesser

**Funding acquisition**

KK Chau, DW Crook, DS Read, N Stoesser, AS Walker
